# Distinct gene expression patterns in vector-residing *Leishmania infantum* identify parasite stage-enriched markers

**DOI:** 10.1101/679712

**Authors:** Iliano V. Coutinho-Abreu, Tiago D. Serafim, Claudio Meneses, Shaden Kamhawi, Fabiano Oliveira, Jesus G. Valenzuela

## Abstract

Promastigotes of *Leishmania infantum* undergo a series of extracellular developmental stages inside the natural sand fly vector *Lutzomyia longipalpis* to reach the infectious stage, the metacyclic promastigote. There is limited information regarding the expression profile of *L. infantum* developmental stages inside the sand fly vector, and molecular markers that can distinguish the different parasite stages are lacking. We performed RNAseq on unaltered midguts of the sand fly *Lutzomyia longipalpis* after infection with *L. infantum* parasites. RNAseq was carried out at various time points throughout parasite development. Principal component analysis mapped the sequences corresponding to the procyclic, nectomonad, leptomonad or metacyclic promastigote stage into distinct positions, with the procyclic stage being the most divergent population. Transcriptional levels across genes varied on average between 10- to 100-fold. Comparison between procyclic and nectomonad promastigotes resulted in 836 differentially expressed (DE) genes; between nectomonad and leptomonad promastigotes in 113 DE genes; and between leptomonad and metacyclic promastigotes in 302 DE genes. Most of the DE genes do not overlap across stages, highlighting the uniqueness of each stage. Furthermore, the different stages of *Leishmania* parasites exhibited specific transcriptional enrichment across chromosomes. Using the transcriptional signatures exhibited by distinct *Leishmania* stages during their development in the sand fly midgut, we determined the genes predominantly enriched in each stage, identifying multiple stage-specific markers for *L. Infantum.* Leading stage-specific marker candidates include genes encoding a zinc transporter in procyclics, a beta-fructofuranidase in nectomonads, a surface antigen-like protein in leptomonads, and an amastin-like surface protein in metacyclics. Overall, these findings demonstrate the transcriptional plasticity of the *Leishmania* parasite inside the sand fly vector and provide a repertoire of stage-specific markers for further development as molecular tools for epidemiological studies.

## Introduction

*Leishmania* parasites are diploid single cell organisms, bearing between 34-36 chromosomes [1]. In clinical isolates, the *Leishmania* karyotype is very plastic, with striking differences not only between geographic isolates [2], but also in parasites isolated from different organs of identical patients [3, 4]. Differences in aneuploidy but also gene copy number variation (CNV) account for most of gene expression variations between *Leishmania* strains or clinical isolates [2]. These differences are associated with *Leishmania* virulence and drug resistance [3, 5], likely representing an evolutionary adaptation for growth in the sand fly vector and human host [3–5]. Surprisingly, < 70 species-specific genes have been found among *Leishmania* species [1].

*Leishmania* are digenetic parasites, switching between mammalian hosts and sand fly vectors. When taken up in a sand fly blood meal, the amastigote stage of *Leishmania* enlarges and exposes the flagellum, undergoing differentiation to the procyclic stage (0.3 fold flagellum to body length ratio) [6]. This transformation is followed by two rounds of cell division inside the insect gut within the confinement of a newly synthesized peritrophic matrix (PM). As the sand fly PM matures, *Leishmania* procyclics elongate their cell bodies to twice the procyclic size, giving rise to the nectomonad stage (0.9 fold flagellum to body length ratio) [6]. Upon breakdown of the PM, the nectomonads escape to the midgut lumen, with some parasites migrating straight to the cardia where they differentiate into haptomonads and eventually form a parasite plug [6, 7]. The free swimming nectomonads attach to the midgut epithelium and give rise to a form displaying a longer flagellum (1.2-1.9 fold flagellum to body length ratio), the leptomonad [6]. Leptomonads undergo multiple rounds of division moving anteriorly along the thoracic midgut. Afterwards, leptomonads begin to shrink their cell bodies and elongate their flagellum, giving rise to the infective forms, the metacyclic parasites (2.0-2.5 fold flagellum to body length ratio). As the infection matures, the proportion of metacyclics relative to the other stages increases with time reaching as high as 80-90% [6–9].

Similar to other Trypanosomatids, *Leishmania* genes are transcribed as long polycistronic RNAs by RNA polymerase II [10, 11]. Such long RNAs are subsequently processed by trans splicing: addition of a capped splice leader sequence encoded in the kinetoplastid DNA (kDNA) at the 5’ end followed up by cleavage and polyadenylation at the 3’ end of each protein-coding unit [10]. Initial microarray studies have identified a very low differential expression (< 5% DE genes) between two *Leishmania* life stages (amastigotes and promastigotes from culture), highlighting a disconnect between transcription and translation. These findings suggest that the *Leishmania* genome is constitutively transcribed, and the control of gene expression is carried out post-transcriptionally at the level of RNA processing and/or translation [11].

Conversely, high throughput RNA sequencing of *Leishmania* transcriptomes detected gene expression differences between intracellular (human host) and extracellular (vector host) parasite stages [12]. Comparing *Leishmania major* culture promastigotes and murine macrophage amastigotes, or culture promastigotes and human macrophage amastigotes, at least 30% of the genes were differentially expressed (q-value <0.05) [13–15]. Apart from a recent work investigating *L. major* stages inside the *Phlebotomus papatasi* sand fly [16], little is known of molecular markers and RNA differential expression between the *Leishmania* promastigote stages developing in the midgut of the sand fly vector, particularly for *L. infantum* inside its natural sand fly vector, *Lutzomyia longipalpis*. In order to fill this knowledge gap, we performed a comprehensive RNAseq investigation to assess *L. infantum* gene expression in the midgut of *Lutzomyia longipalpis* at six time points corresponding to each developmental stage, from procyclic to infective metacyclic promastigotes. Lastly, we identified candidate genes as stage-specific markers for *L. infantum* that will provide a valuable tool for characterizing *Leishmania* stages in the sand fly vector.

## Results

### Parasite growth and differentiation inside the sand fly midgut

The *Leishmania* parasite undergoes markedly different developmental stages inside the sand fly midgut including procyclics, nectomonads, leptomonads, and the infective stage, the metacyclics. We hypothesized that these different developmental stages express a different pattern of transcriptional expression and that this information may define markers to molecularly distinguish distinct parasite stages. To test this hypothesis, we followed the growth and development of *L. infantum* parasites over time inside the midgut of the sand fly *Lu. Longipalpis*. We dissected the sand fly midguts at the six time points after infection (d2, d4, d6, d8, d12, and d14; Figure 1A; Supplemental figure S1A). The parasite growth in the sand fly midgut followed the expected pattern whereby at day 2 (2d), with blood still in the midgut, procyclic promastigotes were the prevailing parasite stage; at day 4 (4d), after blood egestion, there was a low level of parasites (median, 3,000 parasites) and consisted predominantly of nectomonad promastigotes; at day 6 and 8, parasites have multiplied (median, 16,000 parasites on day 6 and 35,000 on day 8) and mostly leptomonad promastigotes were observed (93% on day 6 and 70% leptomonad on day 8). On days 12 and 14, the predominant parasite stage were metacyclic promastigotes (73% on day 12 and 87% on day 14). By day 14, the parasites reached a median of 126,000 parasites per midgut and consisted predominantly of metacyclics (Figure 1, Supplemental figure S1A and B).

**Figure 1.**
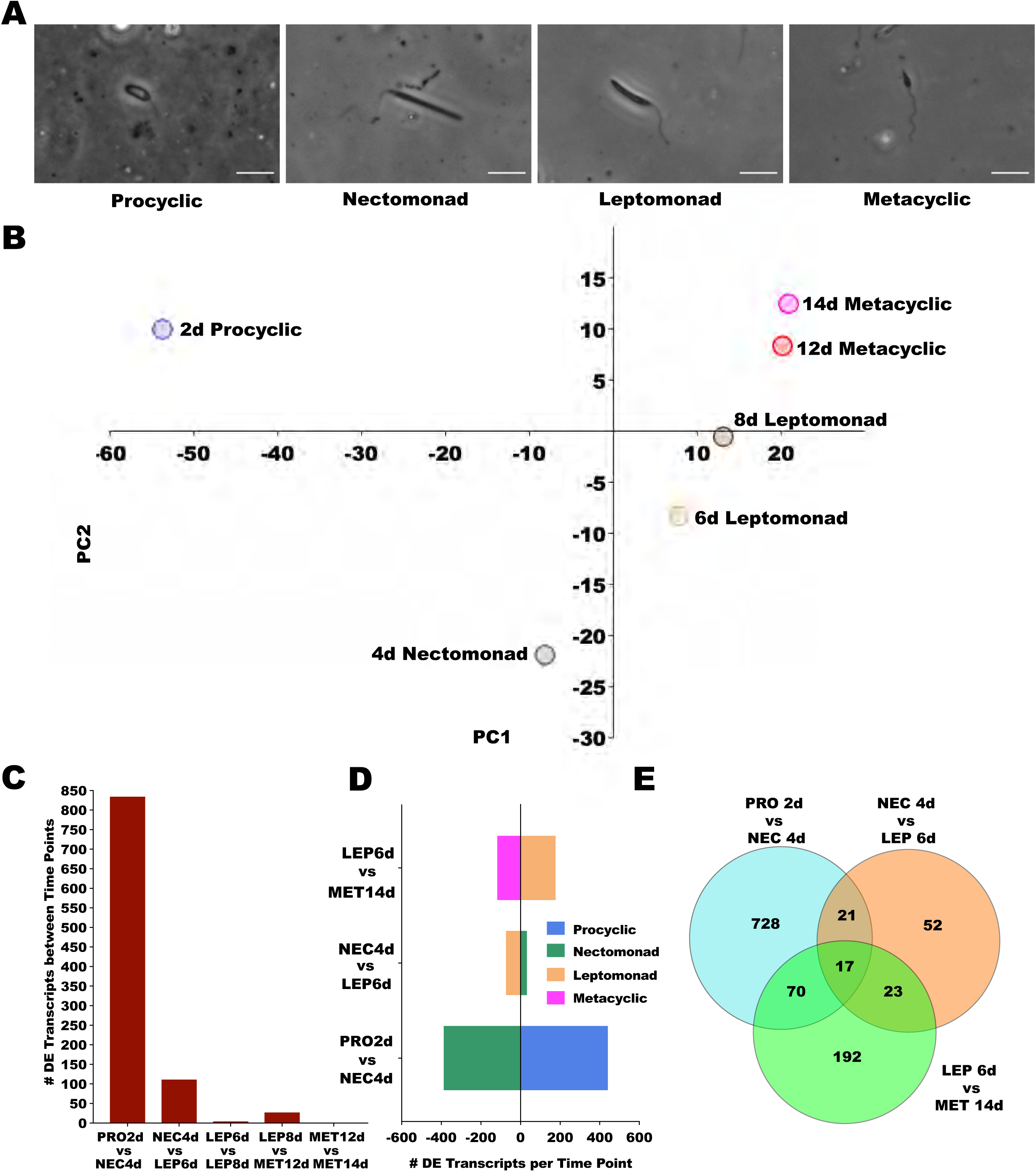
Parasite growth and overall analysis of *Leishmania* sequencing. **A.** Phase contrast images of the *Leishmania* parasites at different stages obtained from midguts at different time points. **B**. Principal component analysis (PCA) describing the position of each *Leishmania* time point in the expression space. Expression space was generated based on the log_2_ TPMs (transcripts per million) of the significantly differentially expressed transcripts across six time points. The Eigenvalues and % variance for PC1 and PC2 were 806.1 and 70.58% and 174.6 and 15.3%, respectively. **C**. Total number of differentially expressed transcripts between *Leishmania* time points. **D**. Enrichment of DE transcripts for each *Leishmania* stage in pairwise comparisons, as color coded in the legend. **E**. Venn diagrams depicting the number of DE transcripts unique and shared amongst pairwise comparisons of *Leishmania* stages. DE was considered significant for transcripts displaying FDR (false discovery rate) q-value lower than 0.05 and LFC (log_2_ fold change) either lower than -0.5 or higher than 0.5. PRO2d: procyclics at day 2. NEC4d: nectomonds at day 4. LEP6d: leptomonads 6 at days. LEP8d: leptomonads at 8 days. MET12d: metacyclics at day 12. MET14d: metacyclics at day 14.

### Gene expression in *Leishmania* residing in the sand fly midgut

We performed RNAseq on RNA extracted from whole *Leishmania*-infected sand fly midguts opting not to purify *Leishmania* parasites to minimize transcriptional noise due to parasite manipulation, but focusing on time points where a specific *Leishmania* stage is predominant to detect differential expression. Experiments were carried out in three biological replicates for d4, d6, d8, d12, and d14, and two biological replicates for d2. All RNAseq libraries gave rise to high quality data and robust expression levels and were used for further analyses.

We mapped the trimmed reads to 8,150 protein-coding genes accounting for all protein-encoding genes identified in the *Leishmania* genome (JPCM5; https://www.sanger.ac.uk/resources/downloads/protozoa/leishmania-infantum.html). Of those, only one third had a known function (Supplemental Fig 2A) represented by categories related to metabolism (met; 21%), signal transduction (st; 18%), protein synthesis machinery (ps; 14%), and protein modification (pm; 11%) (Supplemental Fig S2).

**Figure 2.**
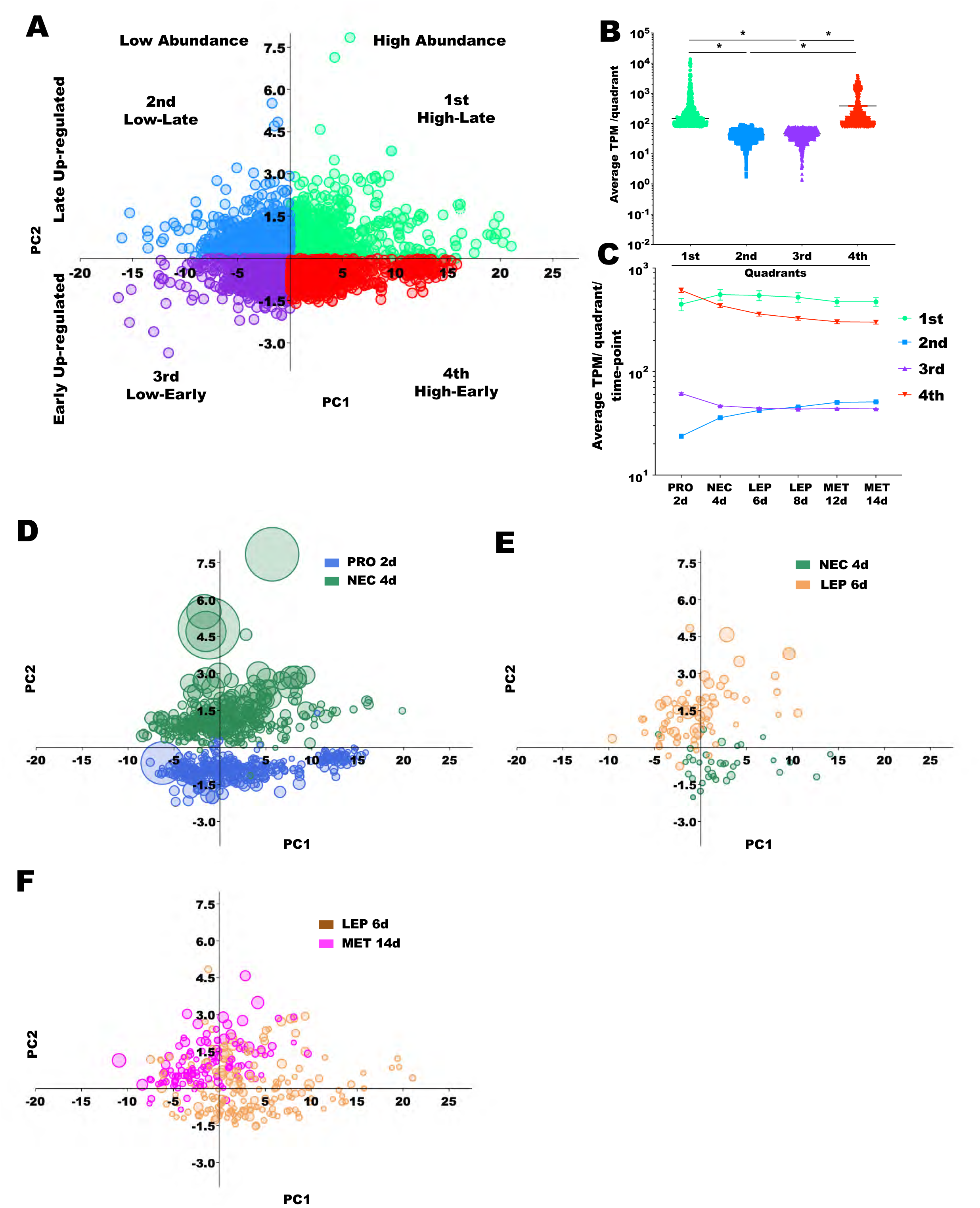
Analysis of differentially expressed (DE) transcript enrichment in different *Leishmania* stages. **A**. Principal Component Analysis (PCA) analysis of all the DE transcripts in all time points based on the log_2_ fold change (LFC) of every pairwise combination of *Leishmania* time points. Each quadrant in the expression space was label from 1^st^ to 4^th^ and the transcripts mapped to the respective quadrants were color coded in Spring Green (1^st^), Dodge Blue (2^nd^), Blue Violet (3^rd^), and Red (4^th^). The Eigenvalues and % variance for PC1 and PC2 % were 20.69 and 95.35% and 0.68 and 3.15%, respectively. HA-LUR: transcripts were high abundance and late up-regulated; LA-LUR: low abundance and late up-regulated genes; LA-EUR: low abundance and early up-regulated genes; HA-EUR: high abundance and early up-regulated genes **B**. Expression analysis per quadrant. The average TPM across time points for every DE transcript mapped in each quadrant was plotted. Horizontal bars indicate median values and differences were statistically significant (* Mann Whitney test, p < 0.0001). Color coding as in A. **C**. Expression analysis per quadrant per time point. The average TPM for each time point for every DE transcript mapped in each quadrant was plotted. Mean TPM as shapes and SEM (standard error of mean) bars are depicted. Based on the differences observed in B and C, the quadrants in A were labeled to describe the up-regulated transcripts expressed in high and low abundance (as defined by PC1) and expressed early and late time points (as defined by PC2). **D-F**. *Leishmania* DE transcripts up-regulated in each stage mapped onto the expression space. D. Bubble plot mapping the procyclic-up-regulated transcripts (Royal blue) and the nectomonad-up-regulated ones (Sea green) on the transcriptional space. E. Bubble plot mapping the nectomonad-up-regulated ones (Sea green) and the leptomonad-up-regulated ones (Saddle brown) on the transcriptional space. F. Bubble plot mapping the leptomonad-up-regulated ones (Saddle brown) and the metacyclic-up-regulated ones (Fuchsia) on the transcriptional space. Differences were statistically significant at p < 0.001 (Chi-square test). DE was considered significant for transcripts displaying FDR q-value lower than 0.05 and LFC either lower than -0.5 or higher than 0.5. PRO2d: procyclics at day 2. NEC4d: nectomonds at day 4. LEP6d: leptomonads at 6 days. MET14d: metacyclics at day 14.

In order to assess the overall similarities in transcriptional profiles amongst *Leishmania* stages, we performed a Principal Component Analysis (PCA) with the overall transcriptional profiles of *Leishmania* at the six time points. This analysis is represented in a two-dimensional plot (Fig 1B; Supplemental fig S3) and summarized in Supplemental Table S1. Interestingly, this unbiased analysis separated the different *Leishmania* stages to the different quadrants of the plot (Fig 1B). As the distance between points correlates with gene expression differences, parasites on day 2 (procyclic stage) were the most divergent population (Fig 1B, left top quadrant). Parasite samples from day 4, which represent nectomonad stage parasites, displayed the second most divergent expression pattern (Fig 1B, left bottom quadrant), followed by day 6 and 8 samples, both enriched with leptomonad-stage parasites (Fig 1B, right bottom quadrant). Parasites on days 12 and 14 mapped closely together (Fig 1B, right top quadrant), indicating very similar gene expression profiles, and, importantly, both of these samples (12 and 14) are enriched in metacylic promastigotes.

The overall pattern of gene expression observed in the PCA of the whole transcriptome (Fig 1B; q-value <0.05; -0.5 < LFC > 0.5) was further analyzed by comparing the differentially expressed (DE) genes between sequential *Leishmania* stages (Fig 1C; Supplemental Table S2). When comparing procyclic stages (2d) and nectomonad stages (4d), we observed 836 differentially expressed genes between these two highly distinct parasite stages (Fig 1C; Supplemental Table S2). When comparing nectomonad (4d) and leptomonad (6d) stages we observed 113 differentially expressed genes (Fig 1C; Supplemental Table S2). Between leptomonad (6d) and metacyclic (14d) stages, 302 genes were differentially expressed (Fig 1C; Supplemental Table S2). As expected, only six genes displayed significant expression differences between leptomonads at day 6 (6d) and leptomonds at day 8 (8d) (Fig 1C) in accordance with their predominance at both timepoints. Along the same lines, there were no differentially expressed genes when comparing metacyclics at day 12 (12d) and metacyclics at day 14 (14d; Fig 1C) suggesting that this parasite stage represents a very homogenous population at these timepoints. Even though a low number of parasites from preceding stages is likely present across most of the studied time points, the predominance of the procyclic stage on 2d, the nectomonad stage on 4d, the leptomonad stage on 6d and 8d, and the metacyclic stage on 12d and 14d (Supplemental fig S1B) was clearly reflected by gene expression differences between time points (Fig 1C).

The pairwise comparisons of DE genes between *Leishmania* stages revealed for the most part an even number of up-regulated genes in each stage (Fig 1D; Supplemental Table S2). When comparing procyclic (2d) and nectomonad (4d) stages, 445 genes were up-regulated in the procyclic stage and 391 genes in the nectomonad stage (Fig 1D, green and blue bars). When comparing nectomonad (4d) versus leptomonad (6d) stages, 36 genes were up-regulated in the nectomonad stage and 77 genes in the leptomonad stage (Fig 1D, light brown and green bars). When comparing leptomonad to metacyclic stages, there were only 181 genes up-regulated in the leptomonad stage (6d) and only 121 genes up-regulated in the metacyclic stage (14d; Fig 1D, light brown and pink bars).

We further determined which genes were DE across multiple stages and those that were DE between only two stages (Fig 1E; Supplemental Table S3). There were 728 DE genes up-regulated in either the procyclic or nectomonad stage, 52 DE genes were up-regulated in either the nectomonad or leptomonad stage, and 192 genes were up-regulated in either the leptomonad or metacyclic stage (Fig 1E; Supplemental Table S4). For the most part, DE genes between two stages were more abundant than DE genes shared by multiple stages, highlighting the existence of transcriptional boundaries for each *Leishmani*a stage (Fig 1E).

We then evaluated whether or not the *Leishmania* genes display different expression patterns throughout development by performing PCA to map the position of all the differentially expressed genes (2,999 differentially expressed genes; q-value <0.05; -0.5 < LFC > 0.5; Supplemental Table S4) in all pairwise comparisons between the different time points onto a two dimensional space (Fig 2A). We observed that DE genes that mapped onto the first quadrant (Fig 2A, top right; Fig 2B, average TPM: 518) and fourth quadrant (Fig 2A, bottom right; Fig 2B, average TPM: 382.5) presented about a ten-fold higher average expression than those DE genes that mapped onto the second quadrant (Fig 2A, top left; Fig 2B, average TPM: 41.99) and third quadrant (Fig 2A, bottom left; Fig 2B, average TPM: 45.76; Supplemental Table S4); therefore, we define the PC1 (top quadrants) as a measure of transcriptional abundance. Further, the differentially expressed genes in the first and second quadrants were up-regulated in late time points, whereas those mapped onto the third and fourth quadrants were up-regulated in the early time points (Figs 2A and C; Supplemental Table S4), suggesting that PC2 accounted for temporal variability in transcription across stages (bottom quadrants). As such, we classify first quadrant transcripts (top right) as high abundance and late up-regulated (High-Late) genes; those that mapped onto the second quadrant (top left) as low abundance and late up-regulated (Low-Late) genes; those that mapped onto the third quadrant as low abundance and early up-regulated (Low-Early) genes, and those that mapped onto the fourth quadrant (bottom right) as high abundance and early up-regulated (High-Early) genes (Figs 2B and C).

Mapping the DE genes enriched in each *Leishmania* stage onto the expression space further underscored the unique expression profiles of the different *Leishmania* stages (Figs 2D-F; Supplemental Fig S3; Supplemental Table S2). When comparing DE genes between the procyclic and nectomonad stages, 98% of the DE genes up-regulated in procyclics mapped onto the Low-Early (43%) and High-Early (55%) quadrants, showing that the procyclic-up-regulated genes were mostly early up-regulated. In contrast, 99% of the nectomonad-up-regulated genes mapped onto the High-Late (50%) and Low-Late (49%; Fig 2D) quadrants showing that the nectomonad-up-regulated genes were predominantly late up-regulated (Chi-square test, p < 0.001; Supplemental Fig S3A). Moreover, there were slightly more DE genes transcribed in high abundance in the procyclic (60%) compared to the nectomonad stage (50%; Fig 2D; Supplemental Fig S3A). When comparing the DE genes in nectomonad and leptomonad stages, we observed that 56% of the nectomonad-up-regulated genes were located on the High-Early, encompassing genes expressed in high abundance and early up-regulated (Fig 2E; Supplemental Fig S3B). Most of the leptomonad-up-regulated genes (59%), on the other hand, belonged to the low abundance/late up-regulated genes by mapping onto the Low-Late quadrant (p < 0.001; Fig 2E; Supplemental Fig S3B). For the DE genes up-regulated in either leptomonad or metacyclic stages (Fig 2F; Supplemental Fig S3C), about 34% of the leptomonad-up-regulated genes mapped onto the High-Late, about one-fourth onto the Low-Late (13%) and Low-Early (14%) quadrants, and the remaining 39% onto the High-Early quadrant. The metacyclic up-regulated genes mapped mostly onto the High-Late (29%) and Low-Late (66%) quadrants (Fig 2F; Supplemental Fig S3C). Hence, the metacyclic up-regulated genes belonged predominantly to the late up-regulated group of genes whereas the leptomonad-up-regulated genes displayed a more broad-spectrum expression pattern (p < 0.001; Fig 2F; Supplemental Fig S3C).

Between-stage differences were also noticed for specific gene families displaying important roles during parasite growth and differentiation in the sand fly vector (Fig 3; Supplemental Table S5). For instance, the number of enriched histone genes, and their expression levels, gradually decreased from the procyclic to the metacyclic stage (Fig 3A; Supplemental Table S5). Genes encoding the small hydrophilic endoplasmic reticulum-associated protein (SHERP) and hydrophilic acylated surface protein a (HASPa), associated to metacylogenesis [17], were up-regulated in leptomonads compared to nectomonads (Fig 3B; Supplemental Table S5), and overall exhibited the highest expression in metacylics (Fig 3B). On the other hand, the gene encoding the META1 protein, also associated to metacylogenesis [17], was up-regulated as early as the nectomonad stage compared to procyclic (Fig 3B; Supplemental Table S5). Regarding the genes involved in the elongation of the glycoconjugate LPG, transcripts for a mannosyltrasferase and the galactosyltransferases (SCG4, SCG7, SCGR3, and SCGR5) were up-regulated in leptomonads and metacyclics, respectively (Fig 3C; Supplemental Table S5). A glycosyltransferase gene (XM_001464195.1), involved in the addition of the LPG’s glucose side chains, was down-regulated in metacyclics (Fig 3C; Supplemental Table S5). Transcription of the gene *ppg4*, responsible for the synthesis of the glycoconjugate proteophosphoglycan, was up-regulated in leptomonads and metacyclics as compared to nectomonads or procyclics (Fig 3C; Supplemental Table S5).

**Figure 3.**
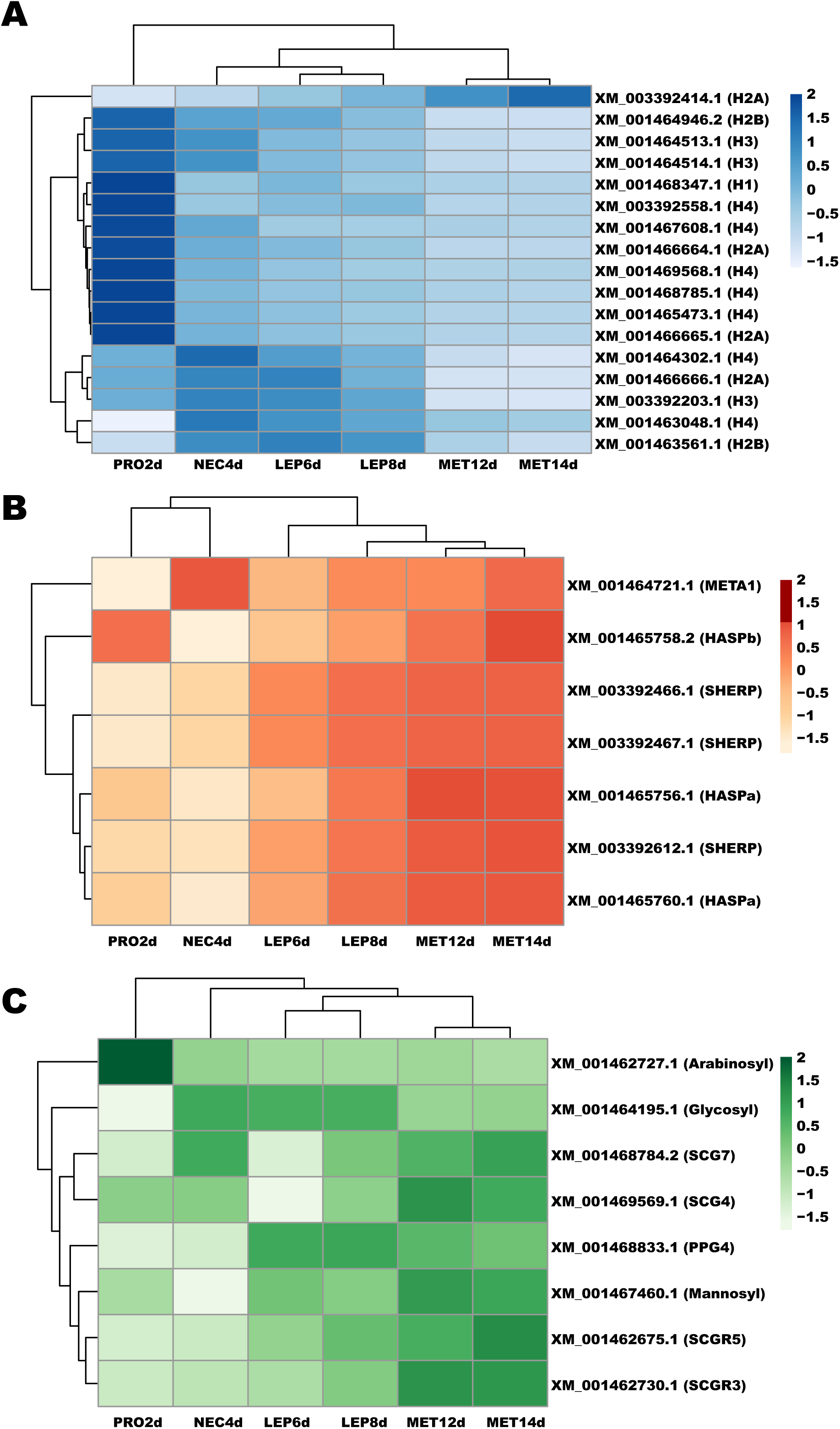
Heatmap depicting temporal expression of selected DE genes. **A.** Genes encoding histone proteins. H1: histone H1. H2A: histone H2A; H2B: histone H2B; H3: histone H3. H4: histone H4. **B.** Metacyclogenesis-related genes. HASPa: hydrophilic acylated surface protein a; HASPb: hydrophilic acylated surface protein b; SHERP: small hydrophilic endoplasmic reticulum-associated protein; META1: META domain-containing protein. **C.** Genes involved in phosphoconjugate sythesis. Arabinosyl: phosphoglycan beta 1,2 arabinosyltransferase; Glycosyl: glycosyltransferase family-like protein; Galactosyl: phosphoglycan beta 1,3 galactosyltransferase; Mannosyl: mannosyltransferase-like protein; PPG4: proteophosphoglycan; LPG3: glucose regulated protein 94. GenBank gene Ids and color intensity scale are also depicted on the left. PRO2d: procyclics at day 2; NEC4d: nectomonads at day 4; LEP6d: leptomonads at day 6; LEP8d: leptomonads at day 8; MET12d: metacyclics at day 12; MET14d: metacyclics at day 14.

Interestingly, the different stages of *Leishmania* parasites exhibited specific transcriptional enrichment across chromosomes (Fig 4; Supplemental Table S6). On chromosome 25, there was a three-fold (or higher) enrichment of genes up-regulated in procyclics compared to nectomonads (Fig 4A). In contrast, three-fold (or higher) enrichment of up-regulated genes in nectomonads was noticed on chromosomes 6, 10, and 31 compared to procyclics (Fig 4B). Between leptomonad and metacyclic, three-fold (or higher) enrichment of up-regulated genes in leptomonads was seen on chromosomes 15, 20, and 33 compared to metacyclics (Fig 4C). On the other hand, five chromosomes (2, 12, 17, 31, and 34) displayed at least three-fold higher abundance of up-regulated genes in metacyclic compared to leptomonads (Fig 4D).

**Figure 4.**
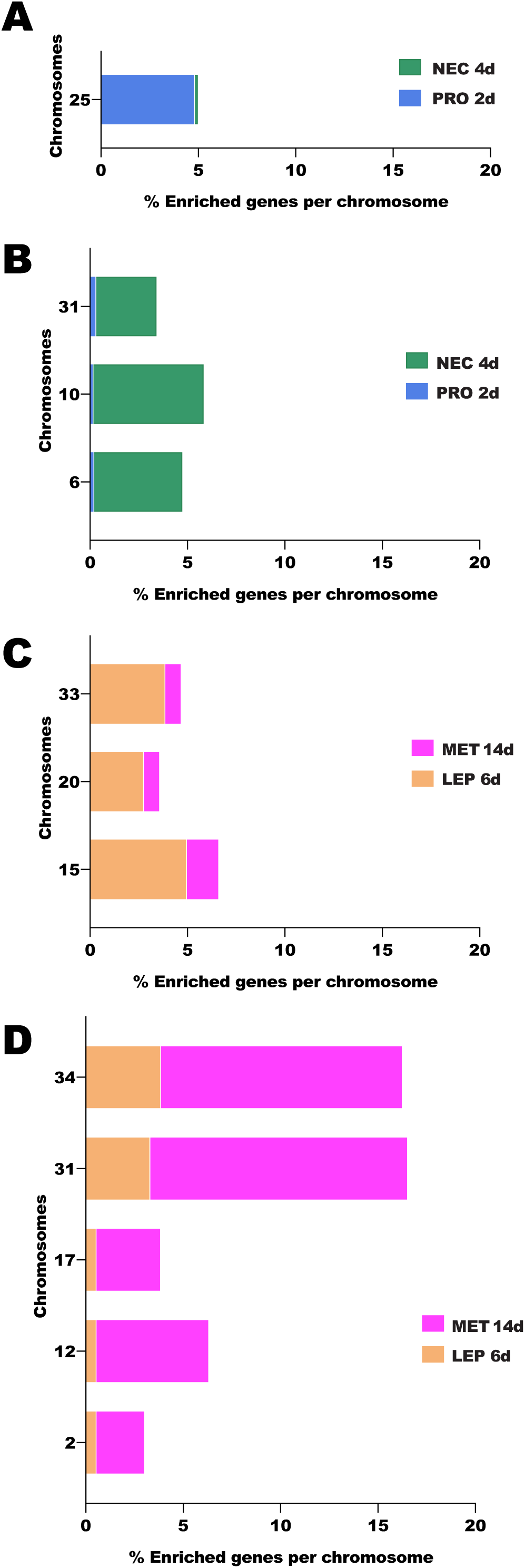
Chromosome displaying at least three-fold enrichment of DE genes across time Leishmania stages. **A.** Chromosomes displaying enrichment of DE genes from procyclic to nectomonad stage. **B.** Chromosomes exhibiting decrease in the proportion of DE genes from procyclic to nectomonad stage. **C.** Chromosomes displaying enrichment of DE genes from leptomonad to metacyclic stage. **B.** Chromosomes exhibiting decrease in the proportion of DE genes from leptomonad to metacyclic stage. PRO2d: procyclics at day 2. NEC4d: nectomonds at day 4. LEP6d: leptomonads at 6 days. MET14d: metacyclics at day 14.

### Candidates for stage-specific molecular markers for *Leishmania infantum*

The differential gene expression between *Leishmania* stages allowed for the identification of stage-specific molecular markers: genes predominantly expressed in one stage compared to all other stages (Fig 5). We initially searched for the genes differentially expressed between one stage and any other stage (Fig 5A-D). For the procyclic, nectomonad, leptomonad, and metacyclic stages, 676, 26, 33, and 175 genes were differentially expressed compared to every other stage, respectively (Fig 5A-D). Among the DE genes between stages, we identified subsets of genes exhibiting stage-specific transcriptional enrichment, i.e. expression levels at LFC > 0.5 and q-value < 0.05 compared to any other stage (Fig 5E; Supplemental Table S7). Among those, 362 genes were up-regulated in procyclics, 5 genes presented a higher expression in nectomonads, 11 genes were up-regulated in leptomonads, and 89 genes displayed metacyclic-specific up-regulation (Fig 5E; Supplemental Tables S8). Among stage-specific candidates, the genes encoding surface proteins are listed in Table 1, with the exception of the nectomonad stage, which was devoid of candidates. For the procyclic stage we identified among other markers the ATPase alpha subunit and the ATP-binding cassette protein subfamily E; for leptomonads we identified among other markers a hypothetical protein similar to a surface antigen-like protein and a putative ATG8/AUT7/APG8/PAZ2; For metacyclics we identified among other markers the surface antigen protein 2, putative amastin-like surface protein, and leishmanolysin. In Table 2, we list the most promising stage-specific makers, encompassing the genes displaying the greatest transcriptional fold change differences to the subsequent stage in development.

**Table 1.** Selected stage specific up-regulated-genes encoding membrane proteins.

| Stage | GeneID | NR best match | Description | e-value | log2 FC<br>PRO2d vs<br>NEC4d | log2 FC<br>PRO2d vs<br>LEP6d | log2 FC<br>PRO2d vs<br>MET14d |
| --- | --- | --- | --- | --- | --- | --- | --- |
| **PRO**<br>**2d** | XM_001462976.2 | XP_001463014.1 | ATPase alpha subunit | 0 | 0.777 | 0.896 | 1.197 |
|  | XM_001465345.1 | XP_001465382.1 | ATP-binding cassette<br>protein subfamily E,<br>member 1 | 0 | 0.854 | 1.243 | 1.430 |
|  |  |  |  |  | log2 FC<br>PRO2d vs<br>LEP6d | log2 FC<br>NEC4d vs<br>LEP6d | log2 FC<br>LEP6d vs<br>MET14d |
| **LEP**<br>**8d** | XM_001462819.1 | XP_001462856.1 | surface antigen-like<br>protein | 0 | -2.862 | -0.888 | 1.435 |
|  | XM_001462818.1 | XP_001462855.1 | surface antigen-like<br>protein | 0 | -2.869 | -0.763 | 1.308 |
|  | XM_001462821.1 | XP_001462858.1 | surface antigen-like<br>protein | 0 | -1.915 | -1.500 | 1.080 |
|  | XM_001463438.1 | XP_001463475.1 | putative<br>ATG8/AUT7/APG8/<br>PAZ2 | 1E-90 | -2.343 | -0.739 | 0.890 |
|  |  |  |  |  | log2 FC<br>PRO2d vs<br>MET14d | log2 FC<br>NEC4d vs<br>MET14d | log2 FC<br>LEP6d vs<br>MET14d |
| **MET**<br>**14d** | XM_001468504.1 | XP_001468541.1 | putative amastin-like<br>surface protein | 1.00E-136 | -1.188 | -1.221 | -0.639 |
|  | XM_001468506.2 | XP_001468543.2 | putative amastin-like<br>surface protein | 0 | -1.090 | -0.958 | -0.705 |
|  | XM_001468449.1 | XP_001468486.1 | putative amastin-like<br>surface protein | 7.00E-139 | -1.694 | -0.842 | -0.719 |
|  | XM_001468507.1 | XP_001468544.1 | putative amastin-like<br>surface protein | 6.00E-138 | -0.957 | -0.974 | -0.798 |
|  | XM_001468501.1 | XP_001468538.1 | putative amastin-like<br>surface protein | 2.00E-149 | -1.476 | -1.135 | -0.893 |
|  | XM_001468436.1 | XP_001468473.1 | putative amastin-like<br>surface protein | 7.00E-123 | -1.696 | -0.801 | -0.659 |
|  | XM_001468503.1 | XP_001468540.1 | putative amastin-like<br>surface protein | 6.00E-136 | -1.681 | -1.651 | -1.211 |
|  | XM_003392619.1 | XP_003392668.1 | putative<br>leishmanolysin | 0 | -2.214 | -1.447 | -0.692 |
|  | XM_001463664.2 | XP_001463701.2 | GP63, leishmanolysin | 0 | -2.896 | -1.470 | -0.707 |
|  | XM_001463660.2 | XP_001463697.2 | GP63, leishmanolysin | 0 | -4.894 | -3.188 | -1.206 |
|  | XM_003392666.1 | XP_003392714.1 | amino acid permease | 0 | -1.586 | -1.535 | -0.870 |
|  | XM_003392664.1 | XP_003392712.1 | amino acid permease | 0 | -2.249 | -2.161 | -0.762 |
|  | XM_001469645.1 | XP_001469682.1 | hypothetical<br>transmembrane<br>protein | 0 | -1.220 | -1.044 | -0.531 |
|  | XM_001462682.1 | XP_001462719.1 | LOW QUALITY<br>PROTEIN: putative<br>ABC1 transporter | 0 | -1.569 | -1.184 | -1.004 |
|  | XM_003392265.1 | XP_003392313.1 | putative surface<br>antigen protein 2 | 2.00E-155 | -2.536 | -1.736 | -1.170 |
|  | XM_001463964.2 | XP_001464001.2 | putative surface<br>antigen protein 2 | 0 | -3.113 | -2.369 | -1.261 |
|  | XM_003392269.1 | XP_003392317.1 | putative surface<br>antigen protein 2 | 0 | -3.771 | -2.919 | -1.498 |
|  | XM_003392274.1 | XP_003392322.1 | surface antigen<br>protein 2 precursor | 0 | -3.653 | -1.558 | -1.105 |

**Table 2.**
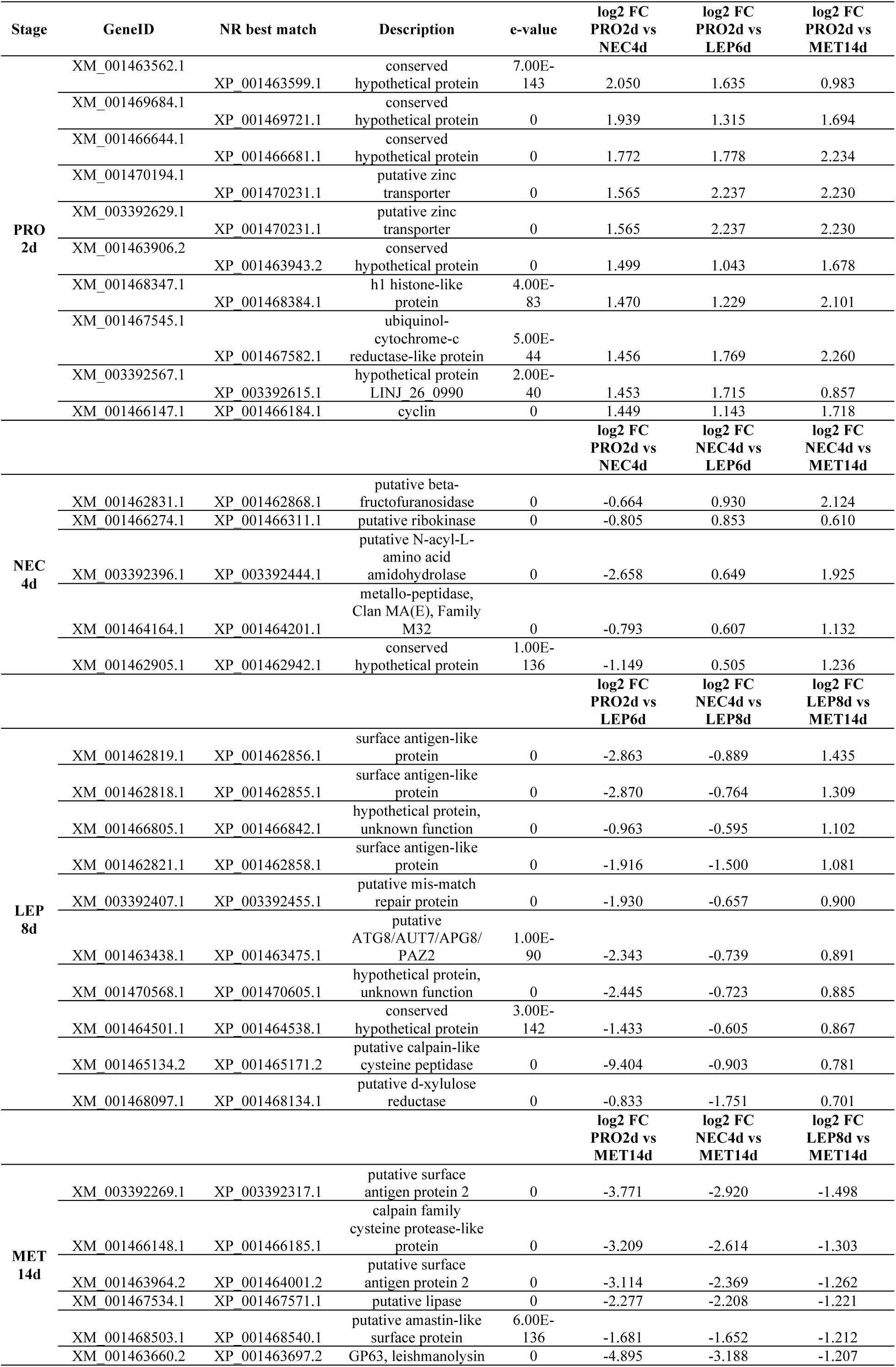
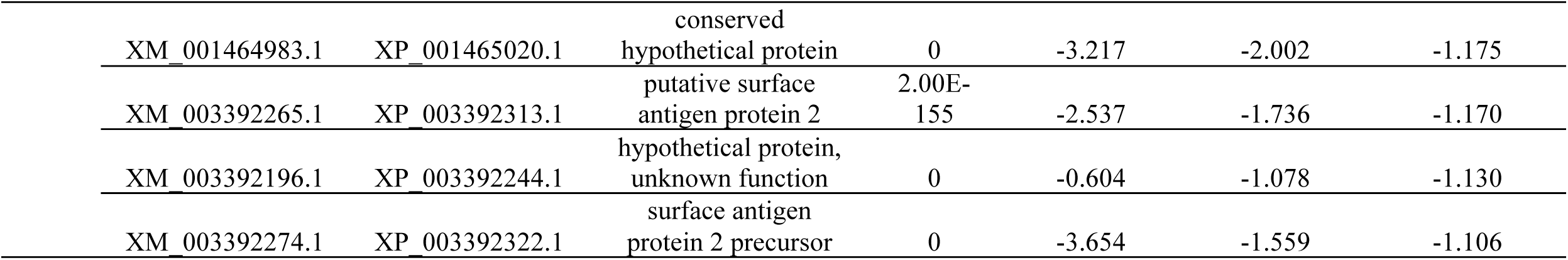
Selected top stage specific markers displaying the highest fold change compared to the next stage.

**Figure 5.**
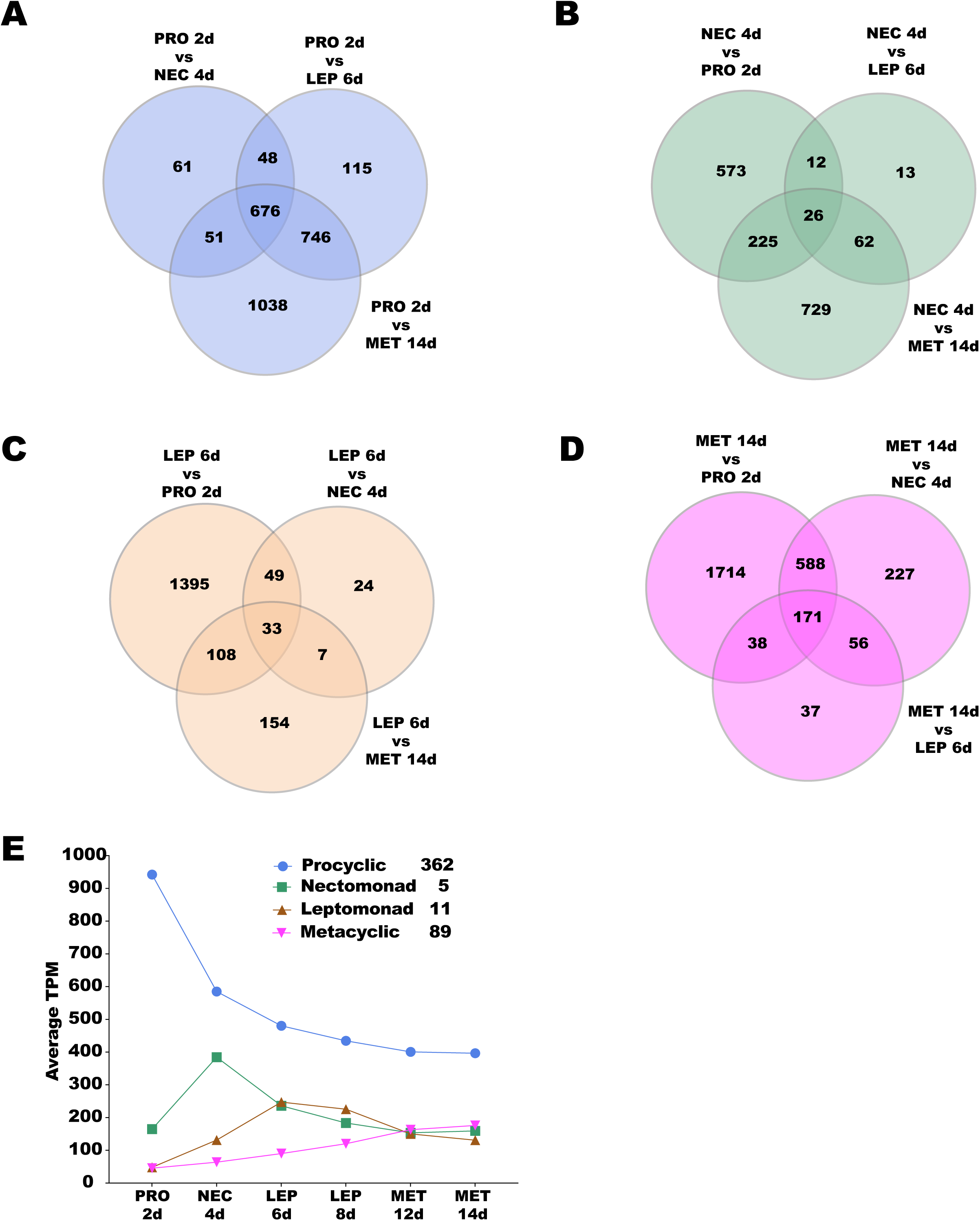
Candidate *Leishmania* stage-specific markers. Venn diagrams highlighting (in white) the numbers of DE genes between (**A**) procyclics, (**B**) nectomonad, (**C**) leptomonad, and (**D**) metacyclic. **E.** Overall expression profile patterns of the candidate *Leishmania* stage-specific markers. Number of candidate genes per stage are shown in the inset. PRO2d: procyclics at day 2. NEC4d: nectomonds at day 4. LEP6d: leptomonads 6 at days. LEP8d: leptomonads at 8 days. MET12d: metacyclics at day 12. MET14d: metacyclics at day 14.

## Discussion

In this study, we hypothesized that different *Leishmania* developmental stages inside the sand fly midgut would have a different pattern of transcriptional expression and that this information could help us to define molecular markers for each of these parasite stages. Our results of high-throughput RNA sequencing of the *L. infantum* stages in the midgut of the sand fly *L. longipalpis* clearly defined the transcriptional boundaries between the different *Leishmania* stages as well as identified gene candidates for *Leishmania* stage-specific molecular markers.

Initial microarrays studies have identified < 5% DE genes between *Leishmania* amastigotes and promastigotes from culture, contrasting to 15% differential expression at the proteomic level [11]. Such a disconnect between transcription and translation suggested that the *Leishmania* genome was constitutively transcribed, and that the control of gene expression was carried out post-transcriptionally at the level of RNA processing and/or translation [11]. In the current work, using statistical settings to detect differential gene expression (q-value < 0.05 and - 0.5 < LFC > 0.5), we have identified 2,999 DE genes amongst all the time points analyzed, which represent 36.8% of the genes in the *L. infantum* genome. These high levels of gene expression plasticity indicates that the *Leishmania* stages exhibiting different morphologies in the sand fly midgut undergo stage-specific changes at the RNA level. These numbers are comparable to previous work studying culture promastigotes versus macrophage amastigotes using less stringent DE statistics; [13–15].

Amongst the DE genes between *L. infantum* midgut stages, we identified genes that were expressed at overall high or low abundance, but also we identified genes up-regulated at early compared to later stages. Also, such sets of DE genes were enriched at different proportions in the different stages. Together, these studies point to the existence of gene expression plasticity at the transcriptional level between *Leishmania* stages. Even though different mechanisms may possibly fine-tune transcriptional expression of *Leishmania* genes [10], the extent of differential expression observed in this study cannot be accounted for by alternative RNA processing between stages [18], suggesting a potential control of gene expression at the transcriptional level in *Leishmania* differentiation that will need to be further explored.

The different stages of *L. infantum* promastigotes also displayed chromosome-specific patterns of enrichment or reduction of DE genes. *L. infantum* tetraploid chromosome 31 [1] displayed a gradual increase in the proportion of upregulated genes from the procyclic to nectomonads and from leptomonads to the metacyclic stage. This may be a mechanism to increase gene expression differences between stages as was reported for *L. mexicana* amastigotes, where the expression of genes located on the tetraploid chromosome 30, a homolog to chromosome 31 in *L. infantum*, was enriched [15]. In fact, other *L. infantum* chromosomes presented an increase or a decrease in the proportion of DE genes as *L. infantum* differentiated from one stage to the next. This phenomenon encompassed not only polysomic (chromosomes 6, 17, 25, 31, and 33) but also multiple disomic (chromosomes 2, 10, 12, 15, 20, and 34) chromosomes [1] across *L. infantum* stages, ruling out chromosomal somy level as a determinant of the differences in the proportions of DE genes detected across *Leishmania* stages. It is noteworthy that the genes differentially expressed during the *Leishmania* ontogeny, which must be hardwired, are also housed on the genetically more stable disomic chromosomes less prone to genetic divergence than their aneuploid counterparts [4].

RNAseq analysis of gene expression between *Leishmania* stages also detected a stronger correlation between gene and protein expression, which had previously been neglected by microarray analysis [11]. Multiple *L. infantum* histones have been shown to be down-regulated during metacyclogenesis *in vitro* [19, 20]. Similarly, multiple histone transcripts were consistently down-regulated throughout *L. infantum* differentiation in *L. longipalpis* midguts in this study, and in *L. major* developing in *P. papatasi* [16]. These findings are in line with the observation that histone gene expression decreases in differentiated cells of higher eukaryotes [21].

The major surface glycan– the lipophosphoglycan (LPG) – of *L. infantum* exhibits glucose side chains, which are maintained during metacyclogenesis [22]. The sugar transferase genes are responsible for the backbone elongation and side-chain decoration of LPG during *Leishmania* metacyclogenesis [23, 24]. Consistent with such a pattern, the glycosyltransferase gene is up-regulated in nectomonads, when LPG is present in high abundance on the parasite’s surface. Similarly, we have identified mRNA up-regulation of galactosyl- and mannosyltransferases in the leptomonad and metacyclic stages, consistent with elongation of LPG in the metacyclic stage [25]. A similar phenomenon was observed for genes linked to *Leishmania* differentiation into infective metacyclics, such as SHERP and HASPa [17]. In accordance with the stationary-phase specific expression of such proteins, the correspondent transcripts are up-regulated in the leptomonad and metacyclic stages. Up-regulation of such genes was also observed in *L. major* metacyclics harvested from sand flies [16]. Along the same lines, one of the genes encoding the *Leishmania* proteophosphoglycan, PPG4 [26, 27], is up-regulated in the leptomonad and maintained at similar levels in the metacyclic stage. At these stages, *Leishmania* secretes a proteophosphoglycan-rich plug in the anterior midgut, which allows these parasites to be regurgitated onto the skin upon sand fly feeding [28, 29]. The concordance in transcript and protein expression amongst these genes, contrasting to the post-transcriptional regulation of the paraflagellar Rod gene [30] and the A600-4 gene [31], clearly demonstrate that *Leishmania* stage-specific protein expression relies on both transcriptional and post-transcriptional controls of gene expression.

One of the gaps in *Leishmania* research is the lack of stage-specific molecular markers. By unveiling the transcriptional boundaries between *L. infantum* stages, this study provides a catalogue of candidates for stage-specific molecular markers that can be tested alone or in combination in in-situ hybridization and Real-Time PCR studies. Such markers will allow the identification of different parasite stages from laboratory culture and vectors, which is important in vector competence and epidemiological studies. Amongst the stage-specific markers, some of which encode surface proteins might facilite the development of monoclonal antibodies and purification of different stages for functional studies. Furthermore, finding that genes encoding surface proteins are enriched in different *Leishmania* stages further supports the fact that surface proteins were one of the principal innovations in the evolution of trypanosomatids [32].

## MATERIAL AND METHODS

### *Leishmania* parasites, sand fly blood feeding and infection, and midgut dissection and storage

The strain of *L. infantum* (MCAN/BR/09/52) used in this study was isolated from a spleen of a dog from Natal, Brazil [33]. The amastigotes used for the sand fly infections were harvested from the spleens of Golden Syrian hamsters, as previously described [34]. Frozen amastigotes were washed once in 1X PBS and five million parasites were inoculated into 1mL of heparinized dog blood. *Leishmania*-seeded blood was loaded into a custom-made glass feeder (Chemglass Life Science, CG183570), capped with a chick skin. The glass feeder was kept at 37°C by circulating heated water. The sand fly *L. longipalpis* was allowed to feed for three hours in the dark. As controls, *L. longipalpis* sand flies were also fed on uninfected heparinized dog blood at the same time. After feeding, fully fed females were sorted and given 30% sucrose solution *ad libitum*. Sand flies from both groups were dissected with fine needles and tweezers on a glass slide at days two, four, six, eight, twelve, and fourteen after blood feeding on RNAse Free PBS (1X). Forty to sixty midguts were quickly rinsed in fresh RNAse Free PBS (1X) and stored in RNAlater (Ambion), following manufacturer’s recommendation. We then performed RNAseq on RNA of *Leishmania*-infected sand fly midguts to prevent potential bias in gene expression that can be generated by purifying *Leishmania* before RNAseq [12]. Experiments were were carried out in three biological replicates.

### Parasite load assessment

A few infected sand fly midguts from all dissected time points were also used to measure parasite loads using Neubauer improved chamber (Incyto, DNC-NO1), as described by the manufacturer. Briefly, dissected midguts were individually transferred to 1.7 mL microtubes (Denville Scientific, C2172) containing 30 μL of 1X PBS and homogenized with a disposable pellet mixer and a cordless motor (Kimble, 7495400000). In order to count the fast moving metacyclic stage parasites, formalin was added to the PBS solution to a final concentration of 0.005%. The Neubauer chamber was loaded with 10 μL of the midgut homogenate (or dilutions of such), and parasites were counted under a microscope (Axiostar plus, Zeiss) at 400X magnification. As the parasite loads on day two are very low and the parasites difficult to be detected due to the blood remains, parasites were not counted at this time point.

### RNA extraction and quality control

Total RNA was extracted using the PureLink RNA Mini Kit (Life Technologies, Carlsbad), following the manufacturer’s recommendations. Briefly, excess RNA later was removed by pipetting, and sample homogenization on lysis buffer was also carried out by pipetting samples up and down for about 60 times. Each sample was eluted into 35μL of RNAse free water. Sample concentration was measured by Nanodrop spectrophotometer (Nano Drop Technologies Inc, Wilmingtom; ND-1000), and RNA quality was assessed by Bioanalyzer (Agilent Technologies Inc, Santa Clara, CA; 2100 Bioanalyzer), using the Agilent RNA 6000 Nano kit (Agilent Technologies) and following the manufacturer’s recommendations. Only one out of the forty eight samples displayed RIN (RNA integrity number) value lower than 7 (Replicate 3 - 14d Pi – RIN 6.7).

### RNAseq library preparation and deep sequencing

The RNASeq library preparation and sequencing was performed at the NC State University Genomic Science Laboratory. The RNAseq libraries were constructed using the NEBNext® Ultra™ RNA Library Prep Kit for Illumina (New England Biolabs, Ipswick MA), following manufacture’s recommendation, in order to obtain reads of 125 nucleotides. RNA libraries were sequenced (Single Ended – 125 SE) in three lanes of the HiSeq 2500 (Illumina, San Diego, CA).

### RNA-seq data trimming, mapping, and differential expression analysis

Raw RNA sequences were trimmed with trimmomatic vs. 0.36 [35] in order to remove poor quality sequences and adaptors using parameters: ILLUMINACLIP:2:30:10 LEADING:3 TRAILING:3 SLIDINGWINDOW:4:15 MINLEN:36. After trimming, quality control of FASTQ sequences were assessed with the FASTQC software (Babraham Bioinfomatics, http://www.bioinformatics.babraham.ac.uk/projects/fastqc/). Trimmed reads were then mapped and counts estimated against the *L. infantum* JPCM5 genome (assembly ASM287v2) using the RNA-Seq by Expectation Maximization (RSEM) vs 1.3.0, Bowtie vs 2-2.2.5 and samtools vs 1.2 [36]. Differential expression among timepoints and conditions were analyzed using the R suite by the Bioconductor package DeSeq2 vs 3.8 [37]. Filtering on all mapped gene counts was performed to exclude genes where the sum of counts in all the conditions was inferior to 10 counts. Default parameters were used with DESeq2 including the shrinks log_2_ fold-change (FC) estimated for each tested comparison [37, 38]. A log_2_ FoldChange and its standard error were generated in addition to a P-value (p value) and a P-adj (Adjusted p-value) to account for the false discovery rate. Significant associations were considered when a P-adj was smaller than 5% (p <0.05) and log_2_ fold change larger than 0.5 (+/-). To classify families of genes in categories (Cs: cytoskeleton; Detox: oxidative metabolism/detoxification; Extmat: extracellular matrix; Imm: immunity; Met: metabolism; Ne: nuclear export; Nr: nuclear regulation; Pe: protein export; Pm: protein modification; Prot: proteosome machinery; Ps: protein synthesis machinery; S: secreted protein; St: signal transduction; Storage: storage protein; Te: transposable element; Tf: transcription factor; Tm: transcription machinery; Tr: transporters and channels) the JPCM5 predicted protein database was blasted using blastp. We automated annotation of proteins was based on a vocabulary of nearly 350 words found in matches to databases including Swissprot, Gene Ontology, KOG, Pfam, and SMART, Refseq-invertebrates, and the diptera subset of the GenBank sequences obtained by querying diptera (organism) and retrieving all protein sequences.

### Data and statistical analyses

Principal component analyses (PCA) were performed with either the log_2_ TPMs or log_2_ fold change (LFC) data using the PAST3 software [39]. This software was also used to construct the bubble plots. Statistical analyses were carried out with PAST3 (multiple Mann Whitney U test) and Prism 7 (GraphPad Software Inc; all the other tests). Venn diagram results were obtained with Venny 2.1 (http://bioinfogp.cnb.csic.es/tools/venny/), and heat-maps/cluster analyses were obtained using the ClustVis tool ([40]; https://biit.cs.ut.ee/clustvis/).

## Acknowledgments

We are indebt with Dr. José M.C. Ribeiro for bioinformatic and scientific discussions and for critical review of the manuscript, and with Dr. Ben Krajacich for R software support. We are also thankful to T.R. Wilson and B.G. Bonilla from LMVR, NIAID for sand fly insectary support. This research was supported by the Intramural Research Program of the NIH, National Institute of Allergy and Infectious Diseases.

## Author contribution

I.V.C.A. and T.D.S. designed and performed the experiments. F.O. supervised bioinformatic analysis. I.V.C.A and F.O. analyzed the data. C.M. performed sand fly insectary work. J.G.V., S.K. and F.O. were involved in the design, interpretation and supervision of this study. I.V.C.A wrote the first draft of the manuscript. J.G.V., S.K. and F.O edited the manuscript.

---

**Figure S1.**
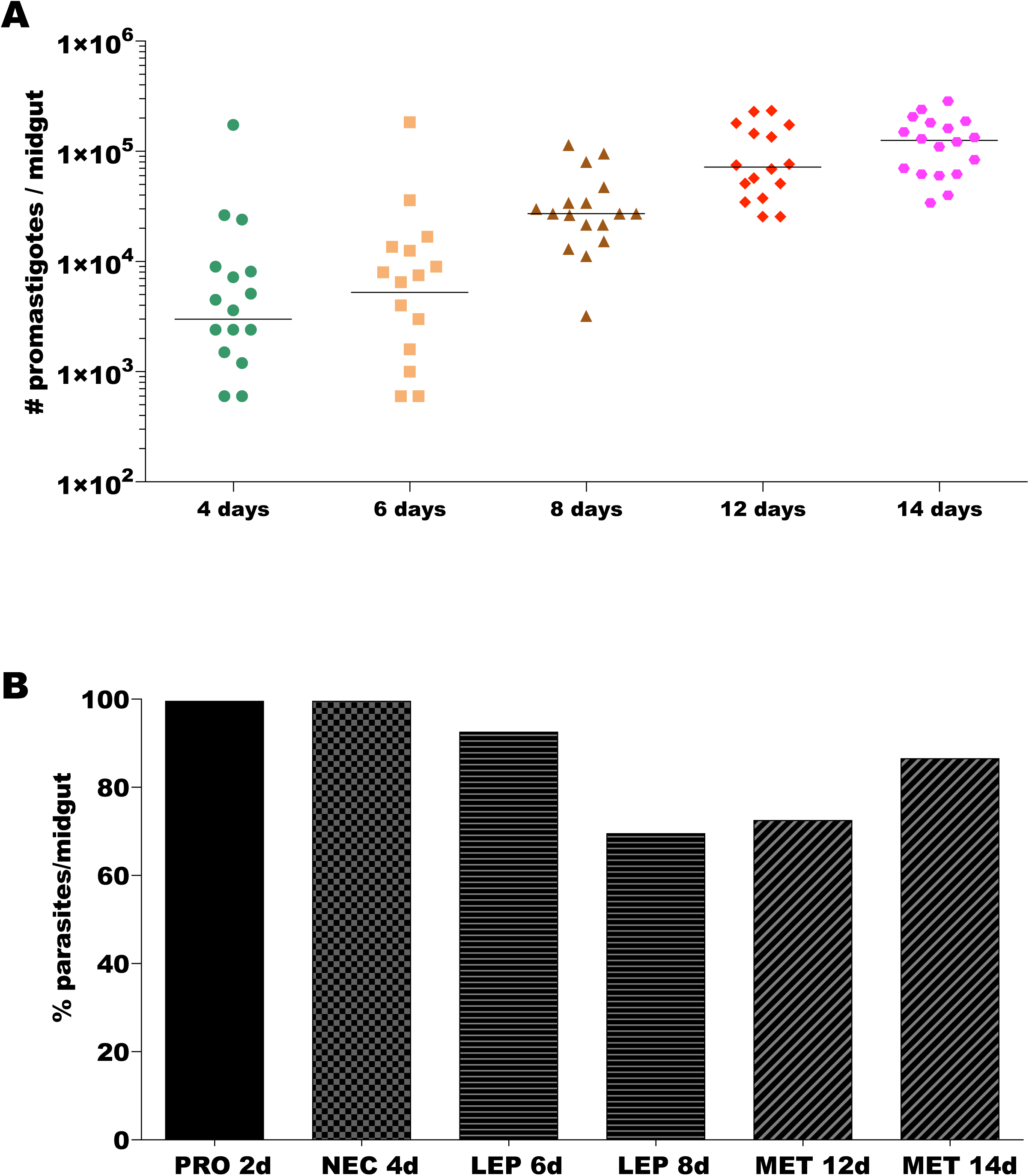
Parasite growth. **A.** Total number of parasites in different time points from single dissected midguts. Horizontal bars indicate median. Pool data from three independent infections. **B**. Proportion of the most predominant Leishmania stage obtained in each time point. PRO2d: procyclics at day 2. NEC4d: nectomonds at day 4. LEP6d: leptomonads 6 at days. LEP8d: leptomonads at 8 days. MET12d: metacyclics at day 12. MET14d: metacyclics at day 14.

**Figure S2.**
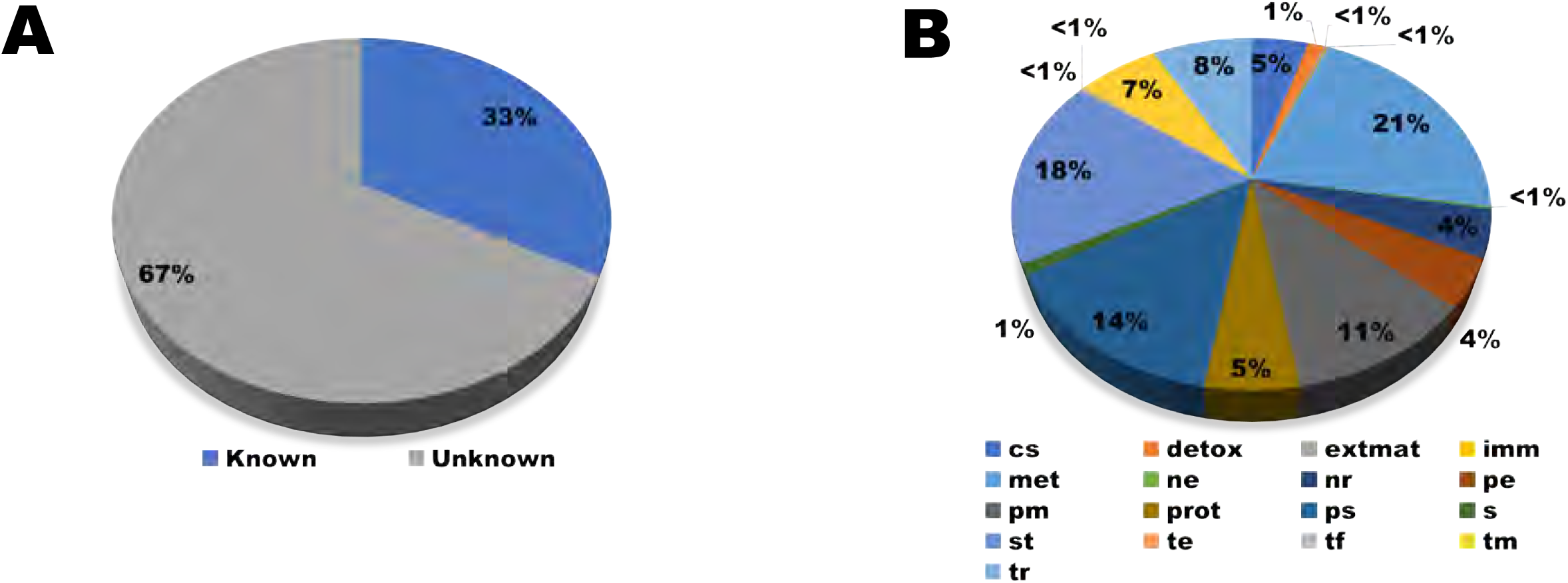
Molecular Functions of the *Leishmania* genes. **A**. Pie chart depicts the overall proportion of transcripts displaying known molecular functions (Known) and orphan sequences (Unknown). **B**. Pie chart displaying the proportion of genes belonging to different molecular functions. Cs: cytoskeleton; Detox: oxidative metabolism/detoxification; Extmat: extracellular matrix; Imm: immunity; Met: metabolism; Ne: nuclear export; Nr: nuclear regulation; Pe: protein export; Pm: protein modification; Prot: proteosome machinery; Ps: protein synthesis machinery; S: secreted protein; St: signal transduction; Storage: storage protein; Te: transposable element; Tf: transcription factor; Tm: transcription machinery; Tr: transporters and channels.

**Figure S3.**
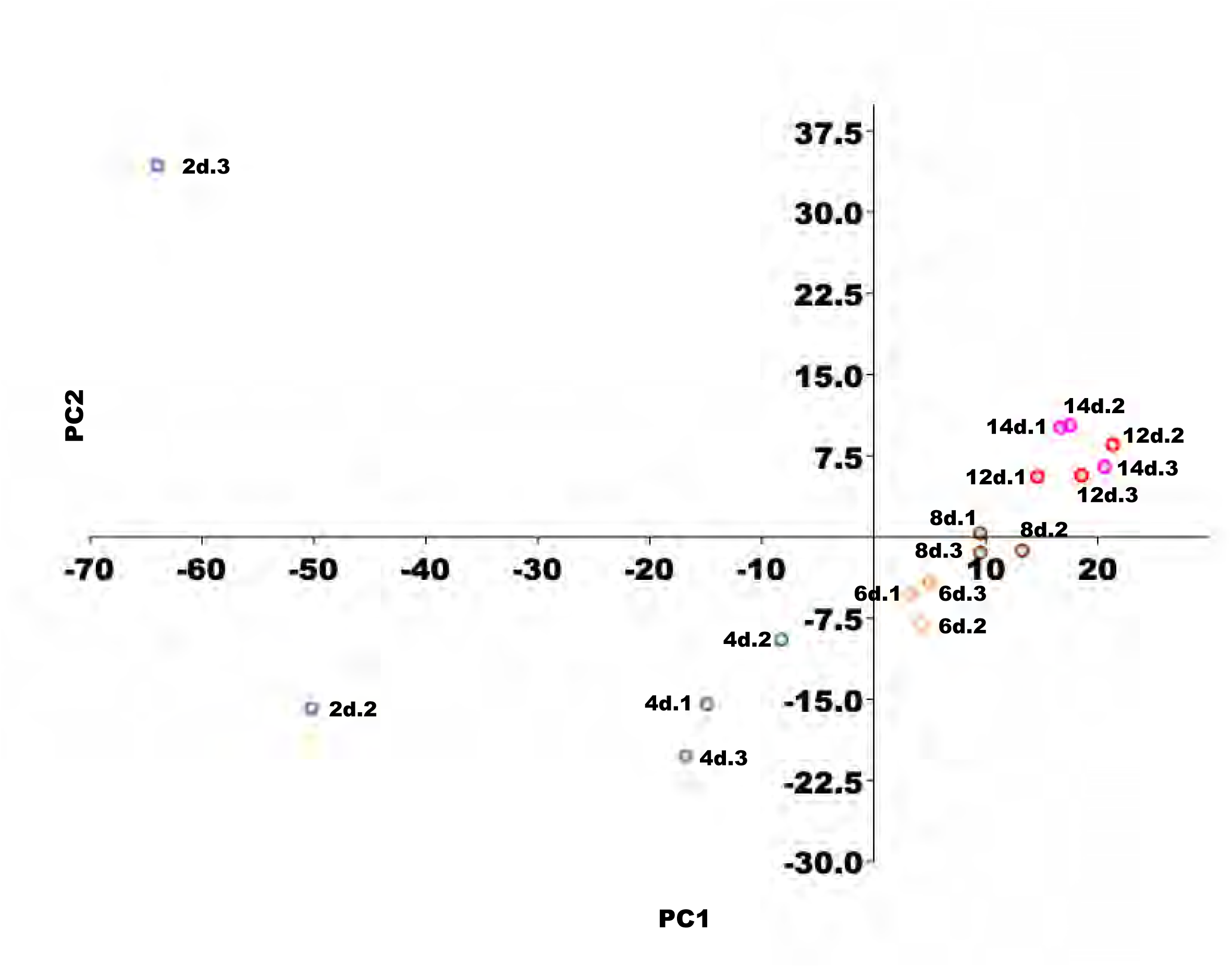
Principal component analysis (PCA) describing the position of each replicate of each *Leishmania* time point in the expression space. Expression space was generated based on the log_2_ TPMs (transcripts per million) using all expressed transcripts across six time points. The Eigenvalues and % variance for PC1 and PC2 were 601.2 and 43.02% and 166.9 and 11.94%, respectively. 2d: procyclics at day 2. 4d: nectomonds at day 4. 6d: leptomonads 6 at days. 8d: leptomonads at 8 days. 12d: metacyclics at day 12. 14d: metacyclics at day 14. Numbers after time points, for instance 2d.2, indicate replicate number.

**Figure S4.**
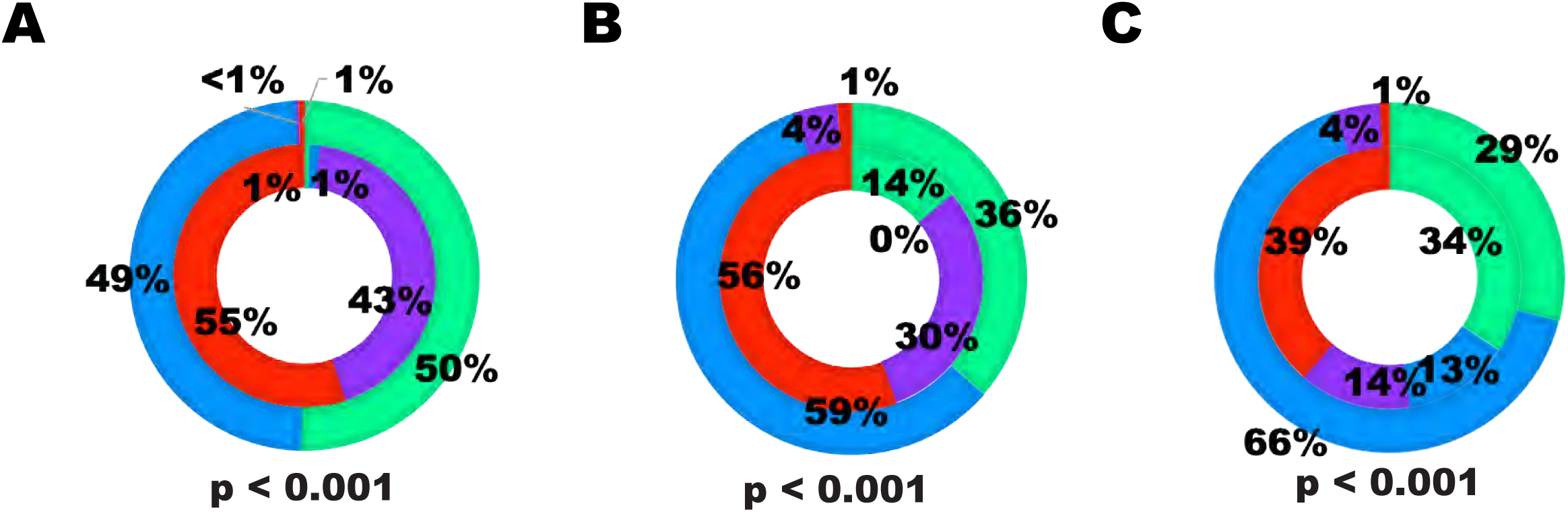
Doughnut chart showing the proportion of enriched transcripts between different stages. (A-C) Procyclic (A), nectomonad (B) and leptomonad (C) stages per quadrant (inner circles) and the proportion of DE transcripts per quadrant in nectomonad (A), leptomonad (B), and metacyclic (C) stages (outer circles). Differences were statistically significant at p < 0.0001 (Chi-square test).

## Supplemental Table Legends

**Table S1. Expression levels (log2 average TPMs per time point) of all genes and PCA coordinates for each time point and for each replicate of each time point.** PCA output: Eigenvalue and % variance. TPM: transcripts per million.

**Table S2. Differential expressed genes between Leishmania stages.** Gene ID number, Principal Component Analysis coordinates, log2 TMP values (transcripts per million), gene annotation information, and counts of TMPs and reads. (PRO2d vs NEC4d) Procyclic over nectomonad comparison. (NEC4d vs LEP6d) Nectomonad over leptomonad comparison. (LEP6d vs MET14d) Leptomonad over metacyclic comparison.

**Table S3. Unique and shared DE genes between stages, as in the Venn diagram (Fig1D).** Gene ID number, log2 Fold change, gene annotation information, and counts of TMPs (transcripts per million) and reads. Positive fold change values indicate enrichment in the former whereas negative fold change values point to enrichment in the later stage.

**Table S4. Overall differential expressed genes across Leishmania stages and DE genes per PCA quadrant.** Gene ID number, Principal Component Analysis coordinates, log2 TMP values (transcripts per million), gene annotation information, and counts of TMPs and reads.

**Table S5. Expression profiles of specific Leishmania genes of known function.** Histones, metacyclogenesis (META1, SHERP, HASPa, HASPb), sugar transferases (glycosyl-, mannosyl-, and galactosyltransferases), and proteophosphoglycan (PPG). Gene ID number, fold change (log2), q-value (padj), gene annotation information (NR best match), e-values, TMP values (log2), average TMPs among replicates. (PRO2d vs NEC4d) Procyclic over nectomonad comparison. (NEC4d vs LEP6d) Nectomonad over leptomonad comparison. (LEP6d vs MET14d) Leptomonad over metacyclic comparison. Regarding fold changes, positive values indicate enrichment in the former whereas negative values point to enrichment in the later stage.

**Table S6. DE genes of each chromosome enriched in each different Leishmania stages**.

**Table S7. DE genes between one Leishmania stage and any other stage.** PRO2d: procyclics at day 2. NEC4d: nectomonds at day 4. LEP6d: leptomonads 6 at days. LEP8d: leptomonads at 8 days. MET12d: metacyclics at day 12. MET14d: metacyclics at day 14.

**Table S8. Candidate markers of each Leishmania stage.** NEC4d: nectomonds at day 4. LEP6d: leptomonads 6 at days. LEP8d: leptomonads at 8 days. MET12d: metacyclics at day 12. MET14d: metacyclics at day 14. TPM: transcripts per million. SD: standard deviation. SEM: standard error of the mean.

## References

1. Rogers MB, Hilley JD, Dickens NJ, Wilkes J, Bates PA, Depledge DP, et al. Chromosome and gene copy number variation allow major structural change between species and strains of *Leishmania*. Genome Res. 2011;21(12):2129–42. Epub 2011/11/01. doi: 10.1101/gr.122945.111. PubMed PMID: 22038252; PubMed Central PMCID: PMCPMC3227102.

2. Iantorno SA, Durrant C, Khan A, Sanders MJ, Beverley SM, Warren WC, et al. Gene Expression in Leishmania Is Regulated Predominantly by Gene Dosage. MBio. 2017;8(5). Epub 2017/09/14. doi: 10.1128/mBio.01393-17. PubMed PMID: 28900023; PubMed Central PMCID: PMCPMC5596349.

3. Zackay A, Cotton JA, Sanders M, Hailu A, Nasereddin A, Warburg A, et al. Genome wide comparison of Ethiopian Leishmania donovani strains reveals differences potentially related to parasite survival. PLoS Genet. 2018;14(1):e1007133. Epub 2018/01/10. doi: 10.1371/journal.pgen.1007133. PubMed PMID: 29315303; PubMed Central PMCID: PMCPMC5777657.

4. Prieto Barja P, Pescher P, Bussotti G, Dumetz F, Imamura H, Kedra D, et al. Haplotype selection as an adaptive mechanism in the protozoan pathogen Leishmania donovani. Nat Ecol Evol. 2017;1(12):1961–9. Epub 2017/11/08. doi: 10.1038/s41559-017-0361-x. PubMed PMID: 29109466.

5. Imamura H, Downing T, Van den Broeck F, Sanders MJ, Rijal S, Sundar S, et al. Evolutionary genomics of epidemic visceral leishmaniasis in the Indian subcontinent. Elife. 2016;5. Epub 2016/03/24. doi: 10.7554/eLife.12613. PubMed PMID: 27003289; PubMed Central PMCID: PMCPMC4811772.

6. Walters LL, Modi GB, Chaplin GL, Tesh RB. Ultrastructural development of Leishmania chagasi in its vector, Lutzomyia longipalpis (Diptera: Psychodidae). Am J Trop Med Hyg. 1989;41(3):295–317. Epub 1989/09/01. PubMed PMID: 2802019.

7. Serafim TD, Coutinho-Abreu IV, Oliveira F, Meneses C, Kamhawi S, Valenzuela JG. Sequential blood meals promote Leishmania replication and reverse metacyclogenesis augmenting vector infectivity. Nat Microbiol. 2018;3(5):548–55. Epub 2018/03/21. doi: 10.1038/s41564-018-0125-7. PubMed PMID: 29556108; PubMed Central PMCID: PMCPMC6007031.

8. Walters LL. Leishmania differentiation in natural and unnatural sand fly hosts. J Eukaryot Microbiol. 1993;40(2):196–206. Epub 1993/03/01. PubMed PMID: 8461893.

9. Lawyer PG, Ngumbi PM, Anjili CO, Odongo SO, Mebrahtu YB, Githure JI, et al. Development of Leishmania major in Phlebotomus duboscqi and Sergentomyia schwetzi (Diptera: Psychodidae). Am J Trop Med Hyg. 1990;43(1):31–43. Epub 1990/07/01. PubMed PMID: 2382763.

10. Clayton CE. Gene expression in Kinetoplastids. Curr Opin Microbiol. 2016;32:46–51. Epub 2016/05/14. doi: 10.1016/j.mib.2016.04.018. PubMed PMID: 27177350.

11. Cohen-Freue G, Holzer TR, Forney JD, McMaster WR. Global gene expression in Leishmania. Int J Parasitol. 2007;37(10):1077–86. Epub 2007/06/19. doi: 10.1016/j.ijpara.2007.04.011. PubMed PMID: 17574557.

12. Patino LH, Ramirez JD. RNA-seq in kinetoplastids: A powerful tool for the understanding of the biology and host-pathogen interactions. Infect Genet Evol. 2017;49:273–82. Epub 2017/02/10. doi: 10.1016/j.meegid.2017.02.003. PubMed PMID: 28179142.

13. Dillon LA, Suresh R, Okrah K, Corrada Bravo H, Mosser DM, El-Sayed NM. Simultaneous transcriptional profiling of Leishmania major and its murine macrophage host cell reveals insights into host-pathogen interactions. BMC Genomics. 2015;16:1108. Epub 2015/12/31. doi: 10.1186/s12864-015-2237-2. PubMed PMID: 26715493; PubMed Central PMCID: PMCPMC4696162.

14. Fernandes MC, Dillon LA, Belew AT, Bravo HC, Mosser DM, El-Sayed NM. Dual Transcriptome Profiling of Leishmania-Infected Human Macrophages Reveals Distinct Reprogramming Signatures. MBio. 2016;7(3). Epub 2016/05/12. doi: 10.1128/mBio.00027-16. PubMed PMID: 27165796; PubMed Central PMCID: PMCPMC4959658.

15. Fiebig M, Kelly S, Gluenz E. Comparative Life Cycle Transcriptomics Revises Leishmania mexicana Genome Annotation and Links a Chromosome Duplication with Parasitism of Vertebrates. PLoS Pathog. 2015;11(10):e1005186. Epub 2015/10/10. doi: 10.1371/journal.ppat.1005186. PubMed PMID: 26452044; PubMed Central PMCID: PMCPMC4599935.

16. Inbar E, Hughitt VK, Dillon LA, Ghosh K, El-Sayed NM, Sacks DL. The Transcriptome of Leishmania major Developmental Stages in Their Natural Sand Fly Vector. MBio. 2017;8(2). Epub 2017/04/06. doi: 10.1128/mBio.00029-17. PubMed PMID: 28377524; PubMed Central PMCID: PMCPMC5380837.

17. Doehl JS, Sadlova J, Aslan H, Pruzinova K, Metangmo S, Votypka J, et al. Leishmania HASP and SHERP Genes Are Required for In Vivo Differentiation, Parasite Transmission and Virulence Attenuation in the Host. PLoS Pathog. 2017;13(1):e1006130. Epub 2017/01/18. doi: 10.1371/journal.ppat.1006130. PubMed PMID: 28095465; PubMed Central PMCID: PMCPMC5271408.

18. Dillon LA, Okrah K, Hughitt VK, Suresh R, Li Y, Fernandes MC, et al. Transcriptomic profiling of gene expression and RNA processing during Leishmania major differentiation. Nucleic Acids Res. 2015;43(14):6799–813. Epub 2015/07/08. doi: 10.1093/nar/gkv656. PubMed PMID: 26150419; PubMed Central PMCID: PMCPMC4538839.

19. Soto M, Iborra S, Quijada L, Folgueira C, Alonso C, Requena JM. Cell-cycle-dependent translation of histone mRNAs is the key control point for regulation of histone biosynthesis in Leishmania infantum. Biochem J. 2004;379(Pt 3):617–25. Epub 2004/02/10. doi: 10.1042/BJ20031522. PubMed PMID: 14766017; PubMed Central PMCID: PMCPMC1224130.

20. Genske JE, Cairns BR, Stack SP, Landfear SM. Structure and regulation of histone H2B mRNAs from Leishmania enriettii. Mol Cell Biol. 1991;11(1):240–9. Epub 1991/01/01. doi: 10.1128/mcb.11.1.240. PubMed PMID: 1986223; PubMed Central PMCID: PMCPMC359614.

21. Stein JL, van Wijnen AJ, Lian JB, Stein GS. Control of cell cycle regulated histone genes during proliferation and differentiation. Int J Obes Relat Metab Disord. 1996;20 Suppl 3:S84–90. Epub 1996/03/01. PubMed PMID: 8680483.

22. Soares RP, Macedo ME, Ropert C, Gontijo NF, Almeida IC, Gazzinelli RT, et al. Leishmania chagasi: lipophosphoglycan characterization and binding to the midgut of the sand fly vector Lutzomyia longipalpis. Mol Biochem Parasitol. 2002;121(2):213–24. Epub 2002/05/30. PubMed PMID: 12034455.

23. Mengeling BJ, Beverley SM, Turco SJ. Designing glycoconjugate biosynthesis for an insidious intent: phosphoglycan assembly in Leishmania parasites. Glycobiology. 1997;7(7):873–80. Epub 1997/11/18. PubMed PMID: 9363429.

24. Novozhilova NM, Bovin NV. Structure, functions, and biosynthesis of glycoconjugates of Leishmania spp. cell surface. Biochemistry (Mosc). 2010;75(6):686–94. Epub 2010/07/20. PubMed PMID: 20636259.

25. Saraiva EM, Pimenta PF, Brodin TN, Rowton E, Modi GB, Sacks DL. Changes in lipophosphoglycan and gene expression associated with the development of Leishmania major in Phlebotomus papatasi. Parasitology. 1995;111 (Pt 3):275–87. Epub 1995/09/01. PubMed PMID: 7567096.

26. Ilg T. Proteophosphoglycans of Leishmania. Parasitol Today. 2000;16(11):489–97. Epub 2000/11/07. PubMed PMID: 11063860.

27. Rogers ME, Ilg T, Nikolaev AV, Ferguson MA, Bates PA. Transmission of cutaneous leishmaniasis by sand flies is enhanced by regurgitation of fPPG. Nature. 2004;430(6998):463–7. Epub 2004/07/23. doi: 10.1038/nature02675. PubMed PMID: 15269771; PubMed Central PMCID: PMCPMC2835460.

28. Rogers ME, Chance ML, Bates PA. The role of promastigote secretory gel in the origin and transmission of the infective stage of Leishmania mexicana by the sandfly Lutzomyia longipalpis. Parasitology. 2002;124(Pt 5):495–507. Epub 2002/06/07. PubMed PMID: 12049412.

29. Rogers ME, Corware K, Muller I, Bates PA. Leishmania infantum proteophosphoglycans regurgitated by the bite of its natural sand fly vector, Lutzomyia longipalpis, promote parasite establishment in mouse skin and skin-distant tissues. Microbes Infect. 2010;12(11):875–9. Epub 2010/06/22. doi: 10.1016/j.micinf.2010.05.014. PubMed PMID: 20561596.

30. Mishra KK, Holzer TR, Moore LL, LeBowitz JH. A negative regulatory element controls mRNA abundance of the Leishmania mexicana Paraflagellar rod gene PFR2. Eukaryot Cell. 2003;2(5):1009–17. Epub 2003/10/14. PubMed PMID: 14555483; PubMed Central PMCID: PMCPMC219351.

31. Murray A, Fu C, Habibi G, McMaster WR. Regions in the 3’ untranslated region confer stage-specific expression to the Leishmania mexicana a600-4 gene. Mol Biochem Parasitol. 2007;153(2):125–32. Epub 2007/04/17. doi: 10.1016/j.molbiopara.2007.02.010. PubMed PMID: 17433460.

32. Jackson AP, Otto TD, Aslett M, Armstrong SD, Bringaud F, Schlacht A, et al. Kinetoplastid Phylogenomics Reveals the Evolutionary Innovations Associated with the Origins of Parasitism. Curr Biol. 2016;26(2):161–72. Epub 2016/01/05. doi: 10.1016/j.cub.2015.11.055. PubMed PMID: 26725202; PubMed Central PMCID: PMCPMC4728078.

33. Aslan H, Dey R, Meneses C, Castrovinci P, Jeronimo SM, Oliva G, et al. A new model of progressive visceral leishmaniasis in hamsters by natural transmission via bites of vector sand flies. J Infect Dis. 2013;207(8):1328–38. Epub 2013/01/05. doi: 10.1093/infdis/jis932. PubMed PMID: 23288926; PubMed Central PMCID: PMCPMC3603531.

34. Gomes R, Teixeira C, Teixeira MJ, Oliveira F, Menezes MJ, Silva C, et al. Immunity to a salivary protein of a sand fly vector protects against the fatal outcome of visceral leishmaniasis in a hamster model. Proc Natl Acad Sci U S A. 2008;105(22):7845–50. Epub 2008/05/30. doi: 10.1073/pnas.0712153105. PubMed PMID: 18509051; PubMed Central PMCID: PMCPMC2397325.

35. Bolger AM, Lohse M, Usadel B. Trimmomatic: a flexible trimmer for Illumina sequence data. Bioinformatics. 2014;30(15):2114–20. Epub 2014/04/04. doi: 10.1093/bioinformatics/btu170. PubMed PMID: 24695404; PubMed Central PMCID: PMCPMC4103590.

36. Li B, Dewey CN. RSEM: accurate transcript quantification from RNA-Seq data with or without a reference genome. BMC Bioinformatics. 2011;12:323. Epub 2011/08/06. doi: 10.1186/1471-2105-12-323. PubMed PMID: 21816040; PubMed Central PMCID: PMCPMC3163565.

37. Love MI, Huber W, Anders S. Moderated estimation of fold change and dispersion for RNA-seq data with DESeq2. Genome Biol. 2014;15(12):550. Epub 2014/12/18. doi: 10.1186/s13059-014-0550-8. PubMed PMID: 25516281; PubMed Central PMCID: PMCPMC4302049.

38. Zhu A, Ibrahim JG, Love MI. Heavy-tailed prior distributions for sequence count data: removing the noise and preserving large differences. Bioinformatics. 2018. Epub 2018/11/06. doi: 10.1093/bioinformatics/bty895. PubMed PMID: 30395178.

39. Hammer O, Harper DAT, Ryan PD. PAST: Paleontological statistics software package for education and data analysis. Palaeontologia Electronica. 2001;4(1):1–9. Epub 22 June 2001.

40. Metsalu T, Vilo J. ClustVis: a web tool for visualizing clustering of multivariate data using Principal Component Analysis and heatmap. Nucleic Acids Res. 2015;43(W1):W566–70. Epub 2015/05/15. doi: 10.1093/nar/gkv468. PubMed PMID: 25969447; PubMed Central PMCID: PMCPMC4489295.

